# Characterization of sterol synthesis in bacteria

**DOI:** 10.1101/047233

**Authors:** Jeremy H. Wei, Xinchi Yin, Paula V. Welander

## Abstract

Sterols are essential components of eukaryotic cells whose biosynthesis and function in eukaryotes has been studied extensively. Sterols are also recognized as the diagenetic precursors of steranes preserved in sedimentary rocks where they can function as geological proxies for eukaryotic organisms and/or aerobic metabolisms and environments. However, production of these lipids is not restricted to the eukaryotic domain as a few bacterial species also synthesize sterols. Phylogenomic studies have identified genes encoding homologs of sterol biosynthesis proteins in the genomes of several additional species, indicating that sterol production may be more widespread in the bacterial domain than previously thought. Although the occurrence of sterol synthesis genes in a genome indicates the potential for sterol production, it provides neither conclusive evidence of sterol synthesis nor information about the composition and abundance of basic and modified sterols that are actually being produced. Here, we coupled bioinformatics with lipid analyses to investigate the scope of bacterial sterol production. We identified oxidosqualene cyclase (Osc), which catalyzes the initial cyclization of oxidosqualene to the basic sterol structure, in 34 bacterial genomes from 5 phyla (Bacteroidetes, Cyanobacteria, Planctomycetes, Proteobacteria and Verrucomicrobia) and in 176 metagenomes. Our data indicate that bacterial sterol synthesis likely occurs in diverse organisms and environments and also provides evidence that there are as yet uncultured groups of bacterial sterol producers. Phylogenetic analysis of bacterial and eukaryotic Osc sequences revealed two potential lineages of the sterol pathway in bacteria indicating a complex evolutionary history of sterol synthesis in this domain. We characterized the lipids produced by Osc-containing bacteria and found that we could generally predict the ability to synthesize sterols. However, predicting the final modified sterol based on our current knowledge of bacterial sterol synthesis was difficult. Some bacteria produced demethylated and saturated sterol products even though they lacked homologs of the eukaryotic proteins required for these modifications emphasizing that several aspects of bacterial sterol synthesis are still completely unknown. It is possible that bacteria have evolved distinct proteins for catalyzing sterol modifications and this could have significant implications for our understanding of the evolutionary history of this ancient biosynthetic pathway.

## Introduction

Sterols are tetracyclic triterpenoid lipids that are required by all eukaryotes for critical cellular functions including maintaining membrane fluidity, phagocytosis, stress tolerance and cell signaling (Bloch, 1991;Swan and Watson, 1998;Castoreno et al., 2005;Xu et al., 2005;Riobo, 2012). Studies on the biosynthesis of sterols in eukaryotes have revealed a variety of novel biochemical reactions while molecular and cell biological studies have revealed unique regulatory mechanisms and key insights into sterol transport (Dimster-Denk and Rine, 1996;Yang, 2006;Nes, 2011). Geochemists also have an interest in these molecules as they have the potential to function as ‘molecular fossils’ (Summons et al., 2006;Love et al., 2009). Sterols, like many polycyclic triterpenoid lipids, are quite recalcitrant and their degradation products, the steranes, are readily preserved in ancient sediments. Sterane signatures in the rock record date as far back as 1.6 billion years (Brocks et al., 2005) and, based on their distribution in modern eukaryotes, are utilized as biomarkers for the existence of specific eukaryotic organisms at the time of deposition (Peters et al., 2007a;b). Because eukaryotes are the predominant extant producers of sterols and because they require sterols for growth, the use of steranes as biomarkers for eukaryotes seems robust. However, sterol production has been observed in a few bacterial species raising the question as to whether bacterial sterol production is significant for the interpretation of sterane signatures (Volkman, 2003;2005).

Bacterial sterol production was first discovered in the aerobic methanotroph *Methylococcus capsulatus* Bath (Bird et al., 1971;Bouvier et al., 1976). *M. capsulatus* was shown to produce 4,4-dimethylcholesta-8,24-dien-3-ol, 4,4-dimethylcholesta-8-en-3-ol, 4-methylcholesta-8,24-dien-3-ol, 4-methylcholesta-8-en-3-ol (Figure 1). Subsequent studies have demonstrated the production of similar sterols in other aerobic methanotrophs of the Methylococcales order within the γ-Proteobacteria (Schouten et al., 2000; Banta et al., 2015). In addition, sterol biosynthesis has also been observed in a few myxobacteria of the δ-Proteobacteria (Bode et al., 2003) and the plantomycete *Gemmata obscuriglobus* (Pearson et al., 2003). *G. obscuriglobus* produces the least biosynthetically complex sterols, lanosterol and the rare lanosterol isomer parkeol. The myxobacteria tend to produce more modified sterols. In particular, *Nannocystis excedens* has been shown to produce primarily cholest-7-en-3-ol (lathosterol) and cholest-8-en-3-ol (Bode et al., 2003).

**Figure 1.**
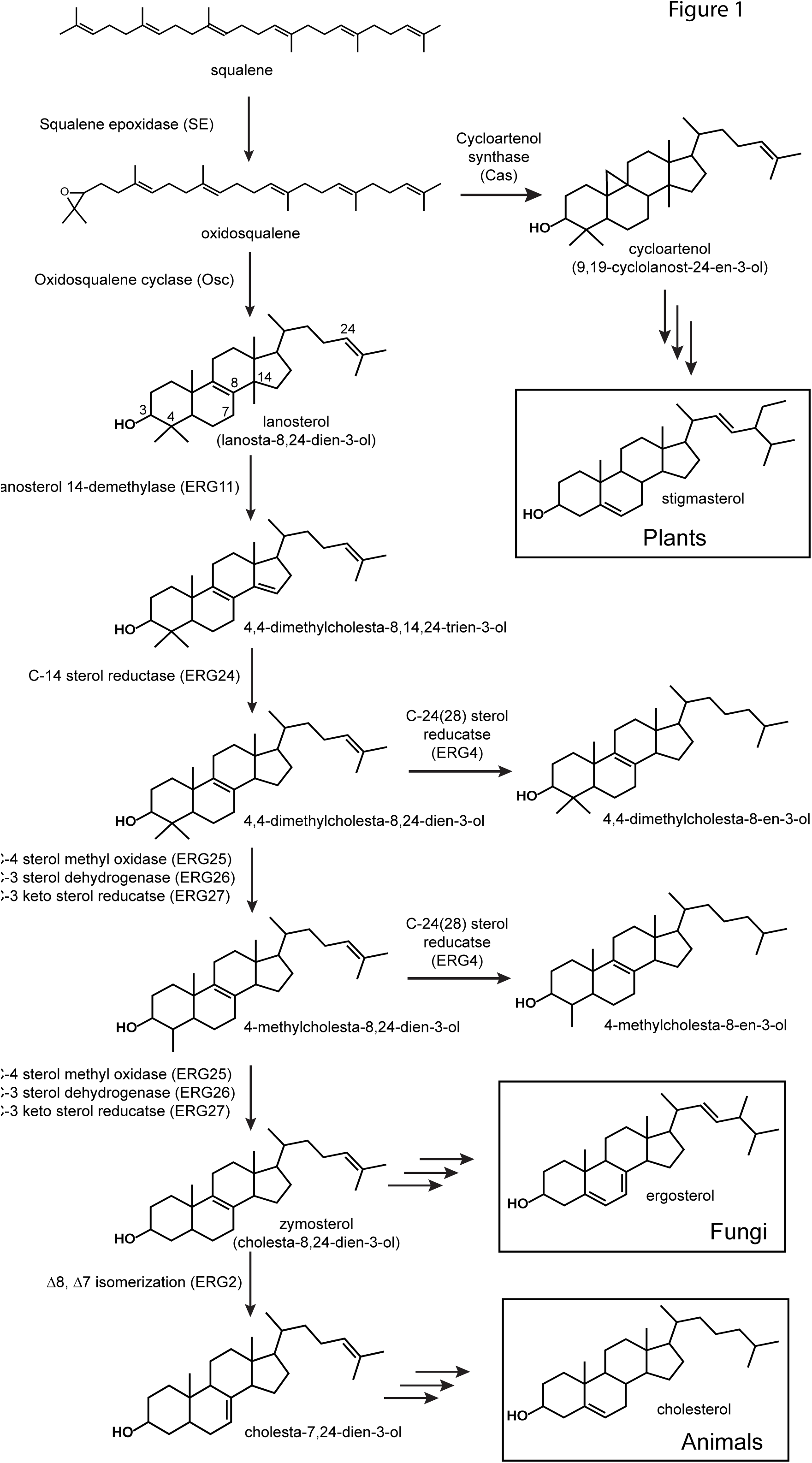
Sterol biosynthesis in eukaryotes. All sterol biosynthetic pathways begin with the oxidation of squalene to oxidosqualene and subsequent cyclization to lanosterol (vertebrates and fungi) or cycloartenol (plants). Shown are the initial enzymatic steps in the conversion of lanosterol to zymosterol which occurs similarly in vertebrates and fungi. Proteins involved in these steps have been characterized from a variety of eukaryotes and the locus to tags shown are those from *Saccharomyces cerevisiae* (Erg).

Recent phylogenetic analyses of bacterial genomes have revealed that the potential for sterol production in this domain might extend beyond what is currently known (Desmond and Gribaldo, 2009;Villanueva et al., 2014). Desmond and Gribaldo identified sterol biosynthesis genes in the myxobacterium *Plesiocystis pacifica* and proposed that this organism had the potential to produce cholesta −7,24-dienol-3β-ol. In addition, Villanueva et al. observed oxidosqualene cyclase (Osc) homologs, required for the initial cyclization of oxidosqualene to lanosterol or cycloartenol (Figure 1) in a variety of bacterial genomes including three aerobic methanotrophs, two Bacteriodetes species and one cyanobacterium symbiont. Further, phylogenetic analyses of bacterial Osc homologs have begun to hint at the evolutionary history of the sterol biosynthetic pathway in bacterial species. In particular, two studies have proposed that bacteria acquired the sterol biosynthetic pathway through horizontal gene transfer from eukaryotes (Desmond and Gribaldo, 2009;Frickey and Kannenberg, 2009).

While genomic and phylogenetic data may provide some clues to the diversity and evolutionary history of the sterol biosynthetic pathway in bacteria, it is important to note that the occurrence of an oxidosqualene cyclase in a bacterial genome demonstrates the potential to produce sterols but it is not conclusive evidence that sterol production is actually occurring. Also, the presence of oxidosqualene cyclase is only indicative of the initial cyclization required to produce the most basic sterols and it does not provide any insight into how sterols may be modified in these Osc containing bacteria. Sterol production is not uniform across all eukaryotes both in terms of the final products produced and in the proteins and enzymatic mechanisms involved in their biosynthesis (Hartmann, 1998;Volkman, 2003;Summons et al., 2006). Vertebrates synthesize cholesterol as a final product while fungi generate ergosterol and plants tend to make stigmasterol (Figure 1). Biosynthetically, plants have a distinct cyclase in which they generate the cyclopropylsterol cycloartenol after cyclization of squalene while vertebrates and fungi generate lanosterol (Figure 1) (Desmond and Gribaldo, 2009). Downstream modifications, including methylations, unsaturation and isomerization, in eukaryotes also differ and it is unclear whether sterol biosynthesis in bacteria is more similar to one (or none) of these pathways.

To fully understand both the potential for sterol production in the bacterial domain and the evolutionary history of the sterol biosynthetic pathway, further studies are needed to characterize sterol production in bacterial species. In this study, we identify potential sterol biosynthesis genes in a variety of bacterial genomes and metagenomes. We then characterized the lipid profiles of a subset of these potential sterol-producers and demonstrate that all but one of the organisms we tested were capable of sterol production under laboratory conditions. Through these studies, it is evident that sterol production is more widespread in the bacterial domain than previously thought and that the bacterial sterol biosynthetic pathway has a complex evolutionary history.

## Materials and Methods

### Bioinformatics Analyses

Homologs of the *Methylococcus capsulatus* Osc protein (locus tag: MCA2873) were identified through BLASTP (Altschul et al., 1997) searches of all bacterial (31,237) and eukaryotic (220) genomes as well as all environmental metagenomes (2,707) on the Joint Genome Institute Integrated Microbial Genomes database (http://img.jgi.doe.gov/). MUSCLE (Edgar, 2004) alignments of bacterial and eukaryotic Osc sequences with an e-value of 1e-50 or lower were generated in Geneious (Biomatters). For metagenomic sequences, only Osc candidates larger than 400 amino acids were included in the alignment. Redundancy in metagenomic alignments was decreased utilizing the Decrease Redundancy web tool (http://web.expasy.org/decrease_redundancy/). Due to large gaps in the metagenomic alignments it was also necessary to extract conserved regions with GBLOCKS (Talavera and Castresana, 2007) prior to building phylogenetic trees containing metagenomics sequences. PhyML (Guindon and Gascuel, 2003) was utilized to generate maximum likelihood phylogenetic trees using the LG+gamma model, four gamma rate categories, ten random starting trees, NNI branch swapping, and substitution parameters estimated from the data. Resulting phylogenetic trees were edited in the Interactive Tree of Life (iTOL) website (http://itol.embl.de/) (Letunic and Bork, 2007;2011).

The following yeast and bacterial proteins were used to search bacterial genomes (BLASTP) for homologs of other sterol biosynthesis proteins: squalene epoxidase (*M. capsulatus* locus tag: MCA2872); lanosterol 14-α-demethylase ERG11 (*Saccharomyces cerevisiae* locus tag: YHR007C); C-14 sterol reductase ERG24 (*S. cerevisiae* locus tag: YNL280C); C-4 methyl sterol oxidase ERG25 (*S. cerevisiae* locus tag: YGR060W); C-3 sterol dehydrogenase ERG26 (*S. cerevisiae* locus tag: YGL001C); 3-keto sterol reductase ERG27 (*S. cerevisiae* locus tag: YLR100W); C-24(28) sterol reductase ERG4 (*S. cerevisiae* locus tag: YGL012W0; 24-dehydrocholesterol reductase (*Homo sapiens* locus tag: HGNC:2859). The e-value cut-off for a potential homolog of these proteins was set at 1e-10 or lower with a minimum 20% identity.

### Lipid Analysis

Bacterial strains surveyed for sterol production and their growth conditions are described in Table 1. All strains were grown in our laboratory except for *Enhygromyxa salina* DSM15201, *Plesiocystis pacifica* SIR-1 DSM14875, and *Sandaracinus amylolyticus* DSM53668. For lipid analysis of these three strains, cells were scraped directly from the agar plates purchased from the German Collection of Microorganisms and Cell Cultures (DSMZ; https://www.dsmz.de/), placed in 2 ml of deionized water and stored at −20°C. All liquid cultures were centrifuged at 5,000 x g for 10 minutes at 4°C and the supernatant was discarded. Cell pellets were frozen at −20°C prior to lipid extraction.

**Table 1.** Bacterial strains tested for sterol biosynthesis

| **Bacteria surveyed in this study** | **Growth conditions** | **Source** |
| --- | --- | --- |
| *Corallococcus coralloides*<br>DSM 2259 | DSMZ Medium 222 at 30°C with shaking at 200 RPM;<br>lipid analysis from 50 ml of a stationary phase culture | DSMZ |
| *Cystobacter fuscus*<br>DSM 2262 | DSMZ Medium 222 at 30°C with shaking at 200 RPM;<br>lipid analysis from 50 ml of a stationary phase culture | DSMZ |
| *Enhygromyxa salina*<br>DSM 15201 | DSMZ Medium 958 on agar plates at 30°C ; lipid<br>analysis from cells scraped off agar plates | DSMZ |
| *Fluviicola taffensis* RW262<br>DSM 16823 | DSMZ Medium 1 for liquid at 30°C with shaking at<br>200 RPM; lipid analysis from 50 ml of a stationary<br>phase culture | DSMZ |
| *Methylobacter luteus* IMV-B-3098 | NMS Medium (Welander and Summons, 2012) plus<br>methane at 5 psi over ambient at 30°C with shaking at<br>200 RPM; lipid analysis from 50 ml of a stationary<br>phase culture | M.G.<br>Kalyuzhnaya,<br>San Diego State<br>University |
| *Methylobacter whittenburyi* ACM<br>3310 | NMS Medium (Welander and Summons, 2012) plus<br>methane at 5 psi over ambient at 30°C with shaking at<br>200 RPM; lipid analysis from 50 ml of a stationary<br>phase culture | M.G.<br>Kalyuzhnaya,<br>San Diego State<br>University |
| *Methyloceanibacter caenitepidi*<br>DSM 27242 | DSMZ Medium 1488 plus 1% methanol at 37°C with<br>shaking at 200 RPM; lipid analysis from 50 ml of a<br>stationary phase culture | M.G.<br>Kalyuzhnaya,<br>San Diego State<br>University |
| *Methylococcus capsulatus* Texas<br>ATCC 19069 | NMS Medium (Welander and Summons, 2012) plus<br>methane at 5 psi over ambient at 30°C with shaking at<br>200 RPM; lipid analysis from 50 ml of a stationary<br>phase culture | ATCC |
| *Methylosarcina lacus* LW14 | NMS Medium (Welander and Summons, 2012) plus<br>methane at 5 psi over ambient at 30°C with shaking at<br>200 RPM; lipid analysis from 50 ml of a stationary<br>phase culture | M.G.<br>Kalyuzhnaya,<br>San Diego State<br>University |
| *Plesiocystis pacifica* SIR-1<br>DSM 14875 | DSMZ Medium 958 on agar plates at 30°C; lipid<br>analysis from cells scraped off agar plates | DSMZ |
| *Sandaracinus amylolyticus*<br>DSM 53668 | DSMZ Medium 67 on agar plates at 30°C; lipid<br>analysis from cells scraped off agar plates | DSMZ |

Frozen cell pellets were resuspended in 2 ml of deionized water and transferred to a solvent washed Teflon centrifuge tube. 5 ml of methanol and 2.5 ml of dichloromethane were added and the cell mixture was sonicated for 1 hour. 10 ml of deionized water and 10 ml of dichloromethane were added to samples after sonication, mixed and incubated at −20°C overnight. Samples were centrifuged for 10 minutes at 2800 x g and the organic layer was transferred to a 40 ml baked glass vial. The total lipid extract was evaporated under N2 and derivatized to acetate or trimethylsilyl (TMS) esters. For derivatization, TLEs were treated with 50 ul of pyridine and 50 ul of acetic anhydride to create acetate derivatives, or with 25 ul of pyridne and 25 ul of TMS + 1% N,O-bis(trimethylsilyl)trifluoroacetamide (BSTFA) for trimethylsilyl derivative compounds. Samples were dried under N2 after derivatization and resuspended in 50-200 μl of dichloromethane prior to high temperature gas chromatography-mass spectrometry (GC-MS) analysis (Sessions et al., 2013).

Lipid extracts were separated on an Agilent 7890B Series GC with helium as the carrier gas at a constant flow of 1.0-1.2 ml/min and programmed as follows: 100°C for 2 min, ramp 15°C/min to 320°C and hold 28-30 min. Analyses were done on a DB5-HT column (30 m X 0.25 mm i.d. X 0.1 μm film thickness) or a DB17-HT column (30 m x 0.25 mm i.d. x 0.15 μm film thickness). 2 μl of the sample were injected into a Gerstel-programmable temperature vaporization (PTV) injector, operated in splitless mode at 320°C. The GC was coupled to a 5977A Series MSD with the source at 230°C and operated at 70 eV in EI mode scanning from 50-850 Da in 0.5 s. All lipids were identified based on their retention time and mass spectra (Figure 2) as well as comparison to prepared internal standards and previously published spectra.

**Figure 2.**
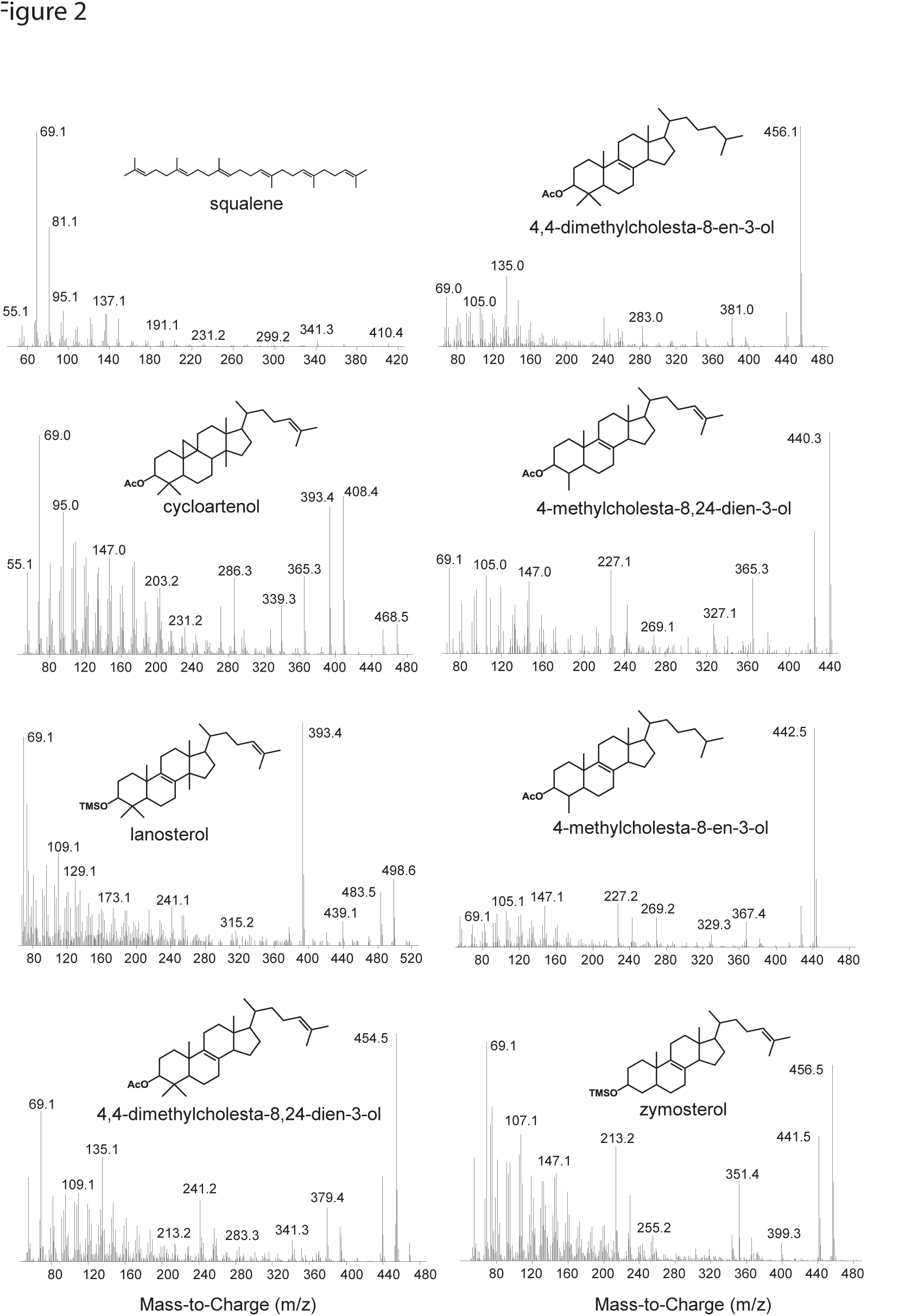
Mass spectra of sterols identified in this study. Spectra of the following acetylated sterols: cycloartenol (9,19-cyclolanost-24-en-3-ol), 4,4-dimethylcholesta-8,24-dien-3-ol, 4,4-dimethylcholesta-8-en-3-ol, 4-methylcholesta-8,24-dien-3-ol, 4-methylcholesta-8-en-3-ol. Spectra of the following trimethylsilylated sterols: lanosterol (lanosta-8,24-dien-3-ol) and zymosterol (cholesta-8,24-dien-3-ol).

## Results

### Identification of oxidosqualene cyclase homologs in bacterial genomes

To identify potential bacterial sterol producers, we queried all bacterial genomes in the Joint Genome Institute Integrated Microbial Genomes database (http://img.jgi.doe.gov/) for homologs of the *M. capsulatus* Bath oxidosqualene cyclase (locus tag: MCA2873). Oxidosqualene cyclases catalyze the conversion of oxidosqualene to lanosterol in vertebrates and fungi and to cycloartenol in land plants (Desmond and Gribaldo, 2009). Deletion of this protein in yeast completely blocks sterol production and therefore its occurrence in a genome is a good indicator of sterol production by an organism (Lees et al., 1995). BLASTP analysis recovered 34 bacterial Osc homologs in five different bacterial phyla with an e-value equal to or lower than e^−100^ and greater than 30% similarity (Table 2). As expected, Osc homologs are found in the genomes of five organisms that have been previously shown to produce sterols: *M. capsulatus* Bath, *N. exedens, Cystobacter fuscus, Stigmatella aurantiaca* and *G. obscuriglobus* (Table 2). The myxobacterium *Corallococcus coralloides* also contains an Osc homolog, however, a previous study of myxobacterial species did not detect any sterols in this bacterium (Bode et al., 2003). Prior phylogenetic studies have also identified Osc homologs in the genomes of *Pleisocystis pacifica* (Desmond and Gribaldo, 2009), *Eudoraea adriatica, Fluviicola taffensis, Methylobacter marinus, Methylomicrobium buryatense* and *Methylomicrobium alcaliphilum* (Villanueva et al., 2014) which we also observed here. However, with the exception of *M. alcaliphilum* (Banta et al., 2015), lipid analysis of these species have not been undertaken to verify sterol production. *Prochloron didenmi* is a cyanobacterial obligate symbiont of the marine ascidian *Lissoclinum patella. P. didenmi* has not been isolated in pure culture but partial genome sequencing of this symbiont has previously revealed an Osc homolog and lanosterol has been observed in whole ascidian extracts (Donia et al., 2011). Our bioinformatics analysis also detected Osc homologs in strains not previously shown to produce sterols including three myxobacteria, ten Methylococcales, two Cyanobacteria, one α-Proteobacterium *(Methyloceanibacter caenitepidi)* and one Verrucomicrobia *(Verrucomicrobiaceae bacterium)*.

**Table 2.**
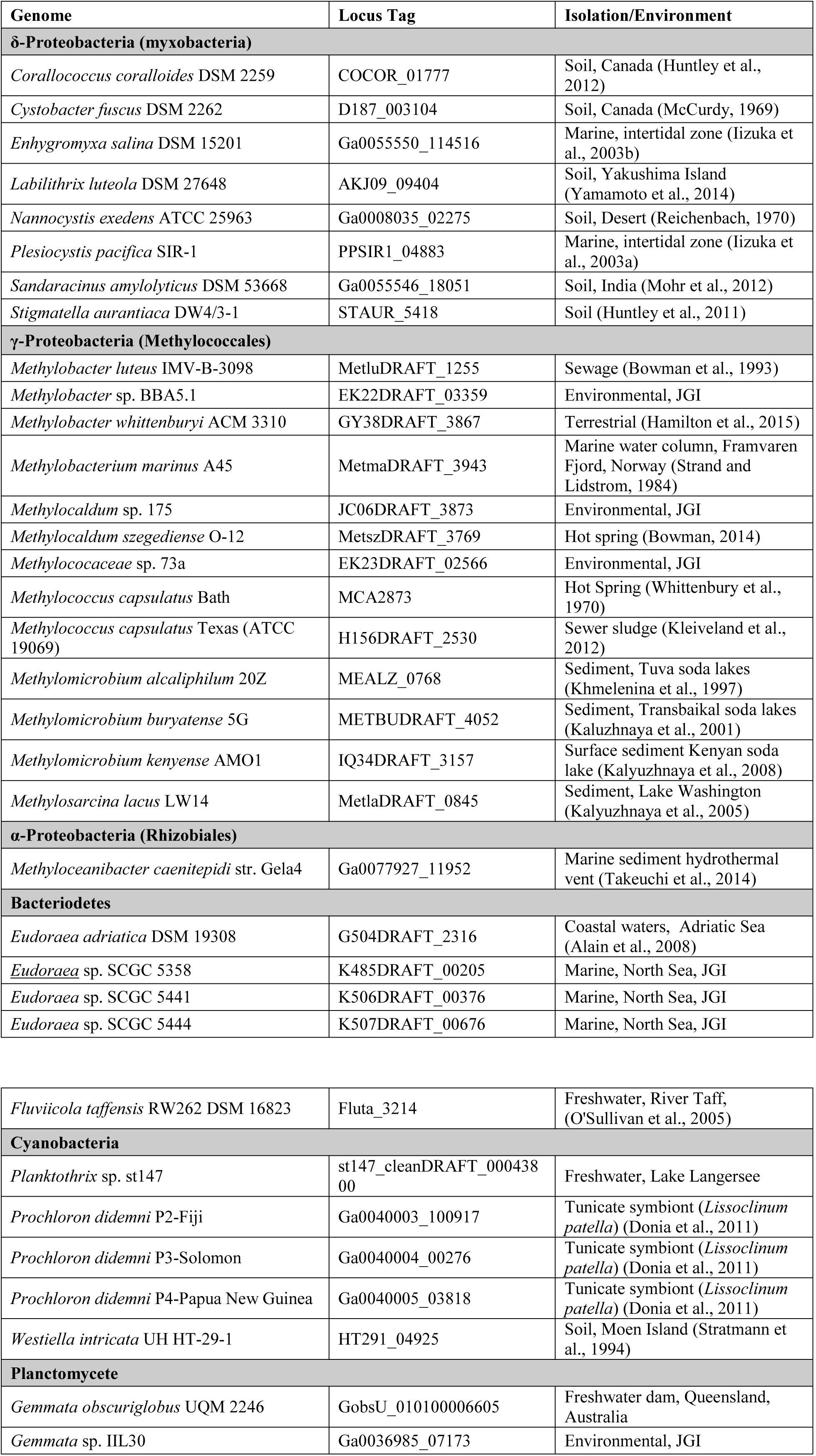
Bacterial genomes that contain oxidosqualene cyclase homologs

| Genome | Locus Tag | Isolation/Environment |
| --- | --- | --- |
| **δ-Proteobacteria (myxobacteria)** |  |  |
| *Coralloccoccus coralloides* DSM 2259 | COCOR_01777 | Soil, Canada (Huntley et al., 2012) |
| *Cystobacter fuscus* DSM 2262 | D187_003104 | Soil, Canada (McCurdy, 1969) |
| *Enhygromyxa salina* DSM 15201 | Ga0055550_114516 | Marine, intertidal zone (Iizuka et al., 2003b) |
| *Labilithrix luteola* DSM 27648 | AKJ09_09404 | Soil, Yakushima Island (Yamamoto et al., 2014) |
| *Nannocystis exedens* ATCC 25963 | Ga0008035_02275 | Soil, Desert (Reichenbach, 1970) |
| *Plesiocystis pacifica* SIR-1 | PPSIR1_04883 | Marine, intertidal zone (Iizuka et al., 2003a) |
| *Sandaracinus amylolyticus* DSM 53668 | Ga0055546_18051 | Soil, India (Mohr et al., 2012) |
| *Stigmatella aurantiaca* DW4/3-1 | STAUR_5418 | Soil (Huntley et al., 2011) |
| **γ-Proteobacteria (Methylococcales)** |  |  |
| *Methylobacter luteus* IMV-B-3098 | MetluDRAFT_1255 | Sewage (Bowman et al., 1993) |
| *Methylobacter* sp. BBA5.1 | EK22DRAFT_03359 | Environmental, JGI |
| *Methylobacter whittenburyi* ACM 3310 | GY38DRAFT_3867 | Terrestrial (Hamilton et al., 2015) |
| *Methylobacterium marinus* A45 | MetmaDRAFT_3943 | Marine water column, Framvaren Fjord, Norway (Strand and Lidstrom, 1984) |
| *Methylocaldum* sp. 175 | JC06DRAFT_3873 | Environmental, JGI |
| *Methylocaldum szegediense* O-12 | MetszDRAFT_3769 | Hot spring (Bowman, 2014) |
| *Methylococaceae* sp. 73a | EK23DRAFT_02566 | Environmental, JGI |
| *Methylococcus capsulatus* Bath | MCA2873 | Hot Spring (Whittenbury et al., 1970) |
| *Methylococcus capsulatus* Texas (ATCC 19069) | H156DRAFT_2530 | Sewer sludge (Kleiveland et al., 2012) |
| *Methylomicrobium alcaliphilum* 20Z | MEALZ_0768 | Sediment, Tuva soda lakes (Khmelenina et al., 1997) |
| *Methylomicrobium buryatense* 5G | METBUDRAFT_4052 | Sediment, Transbaikalian soda lakes (Kaluzhnaya et al., 2001) |
| *Methylomicrobium kenyense* AMO1 | IQ34DRAFT_3157 | Surface sediment Kenyan soda lake (Kalyuzhnaya et al., 2008) |
| *Methylosarcina lacus* LW14 | MetlaDRAFT_0845 | Sediment, Lake Washington (Kalyuzhnaya et al., 2005) |
| **α-Proteobacteria (Rhizobiales)** |  |  |
| *Methyloceanibacter caenitepidi* str. Gela4 | Ga0077927_11952 | Marine sediment hydrothermal vent (Takeuchi et al., 2014) |
| **Bacteroidetes** |  |  |
| *Eudoraea adriatica* DSM 19308 | G504DRAFT_2316 | Coastal waters, Adriatic Sea (Alain et al., 2008) |
| *Eudoraea* sp. SCGC 5358 | K485DRAFT_00205 | Marine, North Sea, JGI |
| *Eudoraea* sp. SCGC 5441 | K506DRAFT_00376 | Marine, North Sea, JGI |
| *Eudoraea* sp. SCGC 5444 | K507DRAFT_00676 | Marine, North Sea, JGI |
| *Fluviicola taffensis* RW262 DSM 16823 | Fluta_3214 | Freshwater, River Taff,<br>(O'Sullivan et al., 2005) |
| **Cyanobacteria** |  |  |
| *Planktothrix* sp. st147 | st147_cleanDRAFT_00043800 | Freshwater, Lake Langersee |
| *Prochloron didemni* P2-Fiji | Ga0040003_100917 | Tunicate symbiont (*Lissoclinum patella*) (Donia et al., 2011) |
| *Prochloron didemni* P3-Solomon | Ga0040004_00276 | Tunicate symbiont (*Lissoclinum patella*) (Donia et al., 2011) |
| *Prochloron didemni* P4-Papua New Guinea | Ga0040005_03818 | Tunicate symbiont (*Lissoclinum patella*) (Donia et al., 2011) |
| *Westiella intricata* UH HT-29-1 | HT291_04925 | Soil, Moen Island (Stratmann et al., 1994) |
| **Planctomycete** |  |  |
| *Gemmata obscuriglobus* UQM 2246 | GobsU_010100006605 | Freshwater dam, Queensland, Australia |
| *Gemmata* sp. IIL30 | Ga0036985_07173 | Environmental, JGI |

A maximum likelihood tree of bacterial and eukaryotic Osc protein homologs was created by aligning the 34 bacterial Osc homologs with 70 eukaryotic Osc sequences and 23 bacterial squalene-hopene cyclase sequences (Shc) as the outgroup. Squalene-hopene cyclases catalyze the conversion of squalene to the polycyclic hopanoid diploptene and are structurally and functionally similar to oxidosqualene cyclases (Siedenburg and Jendrossek, 2011). The phylogenetic tree revealed two bacterial clades of Osc homologs (Figure 3) which is similar to what was observed by Villanueva et al. (Villanueva et al., 2014). One bacterial Osc group (Group 2) falls within the eukaryotic Osc clades while the Group 1 sequences form a sister clade to the eukaryotic cyclases (Figure 3). Further, the clustering of these two bacterial clades of Osc sequences was not congruent with taxonomic phylogeny based on 16S rDNA. For example, we observe Osc homologs from strains of myxobacteria and aerobic methanotrophs clustering in both Group 1 and 2 (Figure 3). This distribution of bacterial Osc homologs suggests a complicated evolutionary history that potentially involves both gene transfer and gene loss.

**Figure 3.**
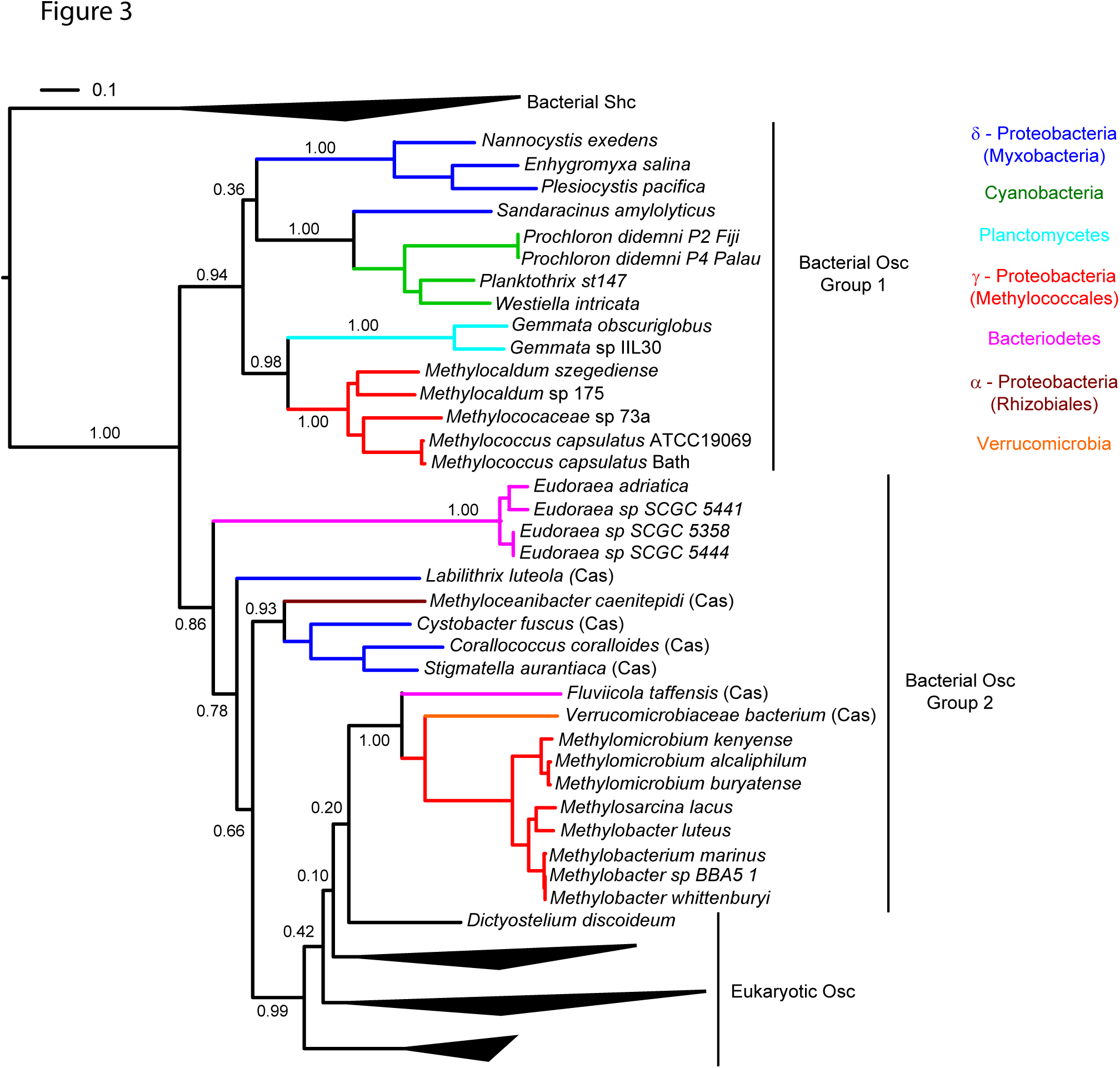
Maximum likelihood phylogenetic tree of oxidosqualene cyclase protein sequences from bacterial and eukaryotic genomes. Bacterial squalene hopene cyclase (Shc) sequences were used as the outgroup. Eukaryotic Osc and bacterial Shc branches are collapsed for better visualization of the tree. Colored branches represent different bacterial phyla: δ-Proteobacteria (blue), Cyanobacteria (green), Planctomycetes (cyan), γ-Proteobacteria (red), Bacteriodetes (pink), α-Proteobacteria (brown) and Verrucomicrobia (orange).

The 34 bacterial species with Osc homologs in their genomes were isolated from a variety of environments indicating that bacterial sterol producers are not restricted to a specific ecological niche (Table 2). The majority of the myxobacterial sterol producers were acquired from soil environments while two other myxobacterial strains originated from marine ecosystems. The Methylococcales species were enriched from a diverse set of ecological settings including sewage sludge, marine water columns, hot springs, freshwater lake sediments and soda lake sediments. Several of the other organisms with Osc homologs in their genomes are also from marine and freshwater environments with *M. caenitepidi* originating from a marine hydrothermal vent, *E. adriatica* from coastal sediments of the Adriatic Sea and *F. taffensis* from sediments of the River Taff.

### Distribution of oxidosqualene sequences in metagenomes

To better understand the ecological distribution of sterol-producing bacteria, we performed a BLASTP search of the environmental metagenomics database on the JGI/ IMG website again utilizing the *M. capsulatus* Bath Osc protein as the query sequence. A total of 176 Osc metagenome sequences were identified with an e-value of e^−50^ or lower. The majority of the metagenomic sequences were from soil, marine or freshwater environments similar to the distribution of isolate environments described above (Figure 4). In addition, Osc sequences were found in metagenomes from estuarine microbial mats, hydrothermal vent fluids and two sequences from sponge symbionts.

**Figure 4.**
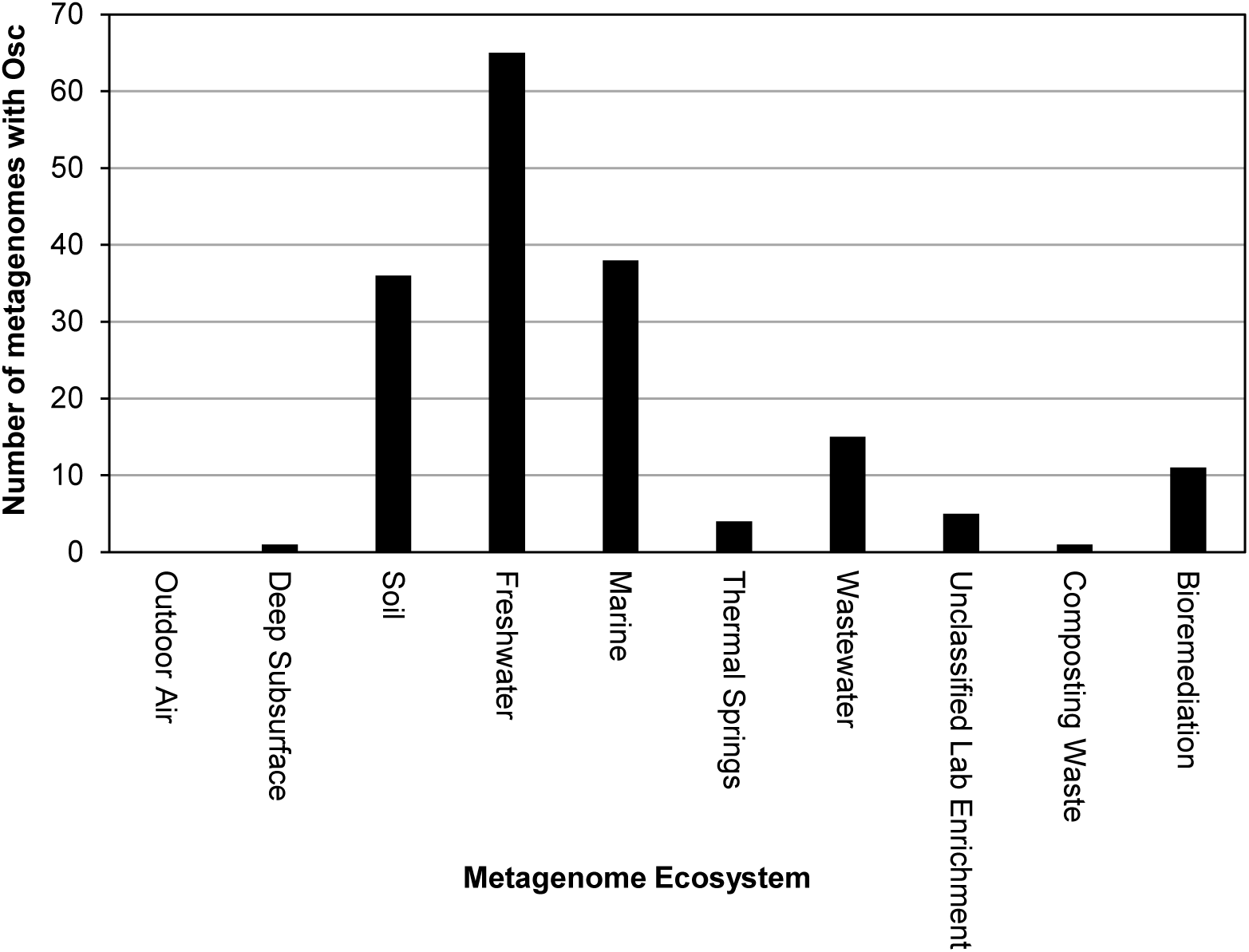
Distribution of Osc protein sequences in metagenomes. Each bar represents the number of Osc homologs identified in the metagenomes from that ecosystem. The majority of homologs were found in freshwater, soil and marine metagenomes.

To distinguish bacterial from eukaryotic Osc sequences in the metagenomes, we generated a maximum likelihood phylogenetic tree of metagenomic and genomic Osc homologs. Because many of the sequences retrieved from metagenomic samples were truncated, we selected the 67 of the 176 metagenomic sequences that were at least 400 amino acids (Osc proteins are generally about 600-650 amino acids) to generate a reliable alignment. After reducing redundancy in the alignment we generated a phylogenetic tree that included a total of 55 metagenomic sequences, 65 eukaryotic genomic sequences and 25 bacterial genomic sequences as well as 18 bacterial squalene-hopene cyclase sequences as the outgroup.

Thirty-seven of the Osc metagenomics sequences retrieved clustered within the two bacterial Osc clades (Figure 5). Some of these sequences grouped with known sterol producers like the Methylococcales and Myxococcales. However, some of these sequences formed their own clades within the bacterial groups or clustered with organisms that have yet to be shown to produce sterols. Thus, identification of these bacterial Osc homologs in metagenome datasets indicates that there are novel sterol-producing bacteria yet to be discovered and that the sterol producing bacteria inhabit diverse environments. Our bioinformatics analysis of metagenomic databases did identify 18 eukaryotic Osc sequences (Figure 6) which were related to algal, plant or fungal Osc homologs. Given how widespread sterol synthesis is in eukaryotes, we had expected to detect more eukaryotic Osc sequences than bacterial sequences in metagenomic databases. However, it has been documented that metagenomic sequencing tends to recover few eukaryotic sequences in general (Lindahl and Kuske, 2013). Therefore, the low number of eukaryotic metagenomic sequences more likely reflects a limited number of eukaryotic sequences in metagenome databases rather than the true prevalence of eukaryotic sterol producers in the environment.

**Figure 5.**
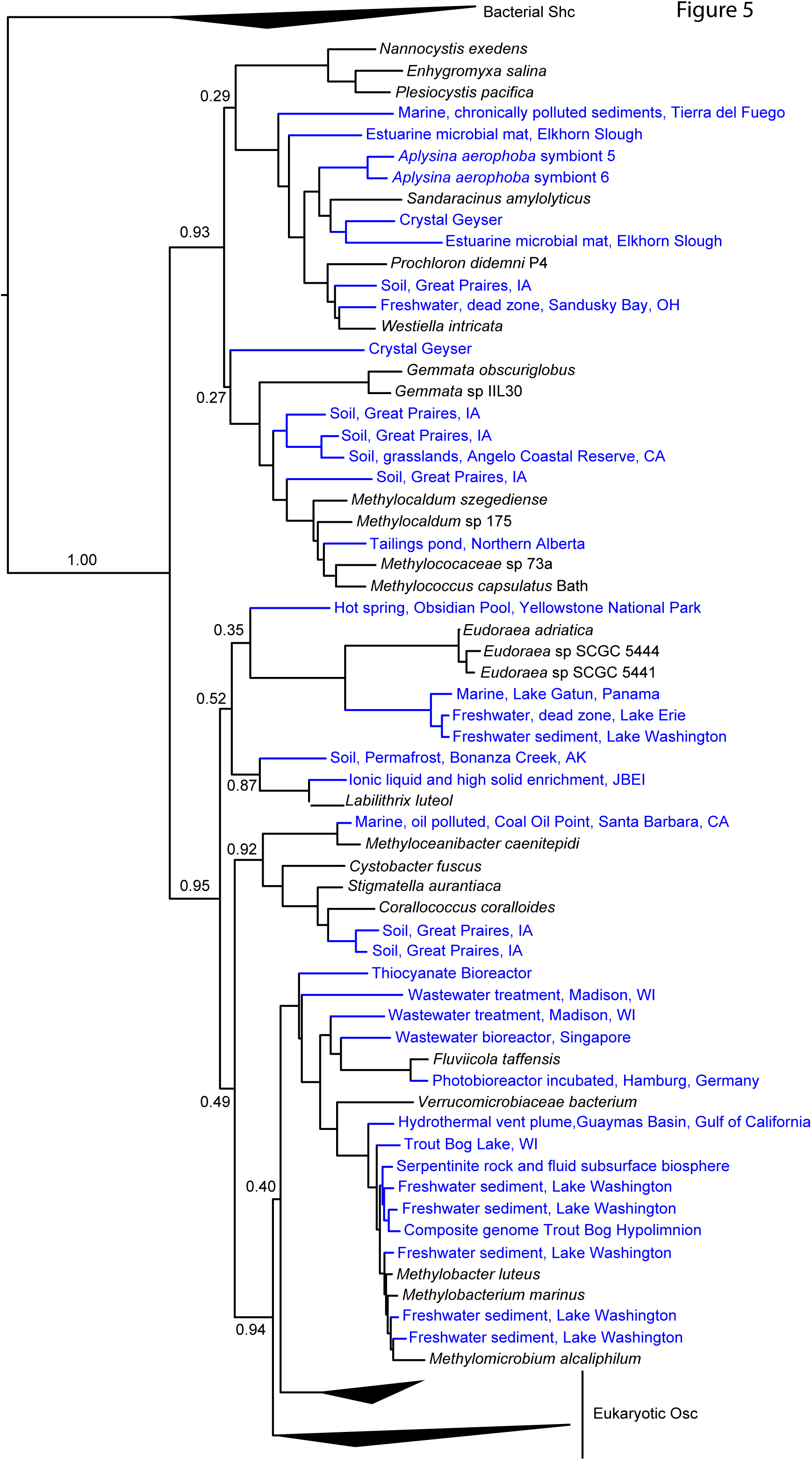
Maximum likelihood phylogenetic tree of bacterial genomic and metagenomic Osc protein sequences. Red branches represent bacterial sequences and black collapsed branches are eukaryotic sequences. Blue labels indicate metagenomic sequences and black labels indicate sequences from genomes. Bacterial Shc sequences were used as the outgroup.

**Figure 6.**
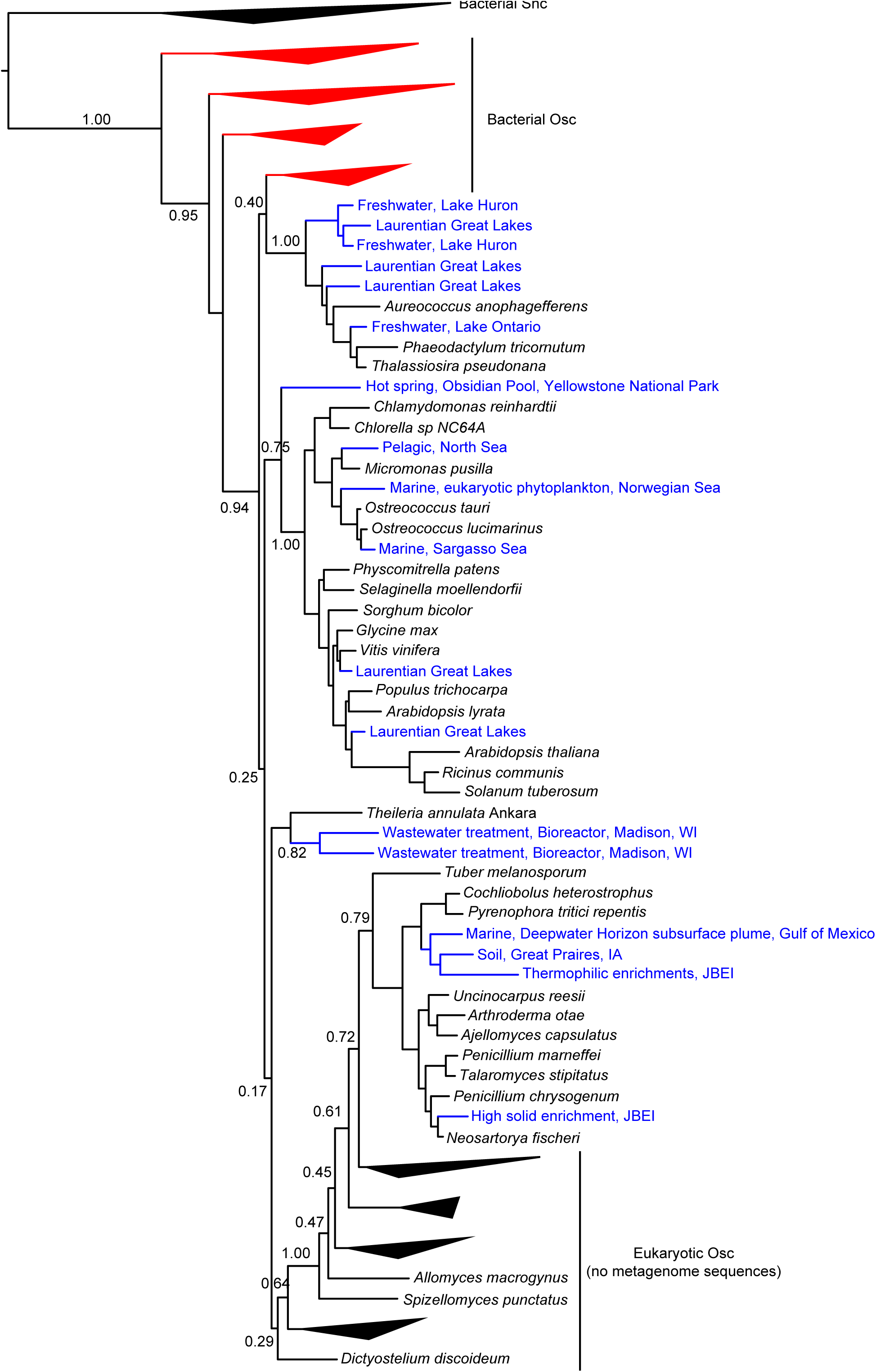
Maximum likelihood phylogenetic tree of eukaryotic genomic and metagenomic Osc protein sequences. Red collapsed branches are bacterial sequences and black branches represent eukaryotic sequences. Blue labels indicate metagenomic sequences and black labels indicate sequences from genomes. Eukaryotic branches with no metagenomic Osc sequences were collapsed to reduce the size of the tree. Bacterial Shc sequences were used as the outgroup.

### Lipid analysis of potential sterol producers

Our identification of Osc homologs in bacterial genomes demonstrates that the potential for sterol synthesis exist in a variety of bacteria. However, the majority of these potential sterol-producing bacterial strains have not been tested for sterol production. In addition, the occurrence of Osc in a genome only suggests the production of the most basic sterols, lanosterol or cycloartenol. Thus, lipid analysis is needed not just to verify sterol production but also to determine if and how sterols are modified in bacteria. We performed lipid analysis on 11 Osc-containing bacteria that included five myxobacteria, four Methylococcales, one Bacteriodetes and one α-Proteobacterium (Table 1). In addition, we searched the genomes of these 11 organisms for other sterol biosynthesis protein homologs. Our goal was to link the occurrence of these downstream biosynthesis genes with any sterol modifications, such as saturations and demethylations, these bacteria may be carrying out.

### Sterol production in the myxobacteria

Four of the five myxobacterial strains tested were found to produce sterols (Table 3). *C. fuscus* strains were previously reported to produce either lanosterol or cycloartenol (Bode et al., 2003) and the *C. fuscus* strain we analyzed produced cycloartenol. We identified homologs for C-14 demethylation and C-24 reduction in the *C. fuscus* genome but did not observe any sterols with these modifications (Table 4). The other three myxobacteria, *E. salina, P. pacifica* and *S. amylolyticus* all produced lanosterol rather than cycloartenol and all three strains modified lanosterol to generate zymosterol (cholesta-8,24-dien-3-ol) (Figure 1 and Figure 7). The conversion of lanosterol to zymosterol requires demethylation at C-4 and C-14 and a reduction at C-14. A previous study had identified homologs for these biosynthetic steps in the *P. pacifica* genome (Desmond and Gribaldo, 2009) and we also observe these protein homologs in *E. salina* and *S. amylolyticus*. However, C-4 demethylation requires three proteins in yeast and vertebrates (ERG25, ERG26 and ERG27) and we do not observe homologs to all of these proteins in these three strains. Desmond et al. pointed out that *P. pacifica* did not have a homolog for one of the three proteins required for C-4 demethylation (ERG27) and we demonstrate that *S. amylolyticus* is also missing this protein. Further, we were only able to identify one homolog of these three proteins in *E. salina* (Table 4). Thus, it is unclear how these myxobacterial strains are fully demethylating at the C-4 position.

**Figure 7.**
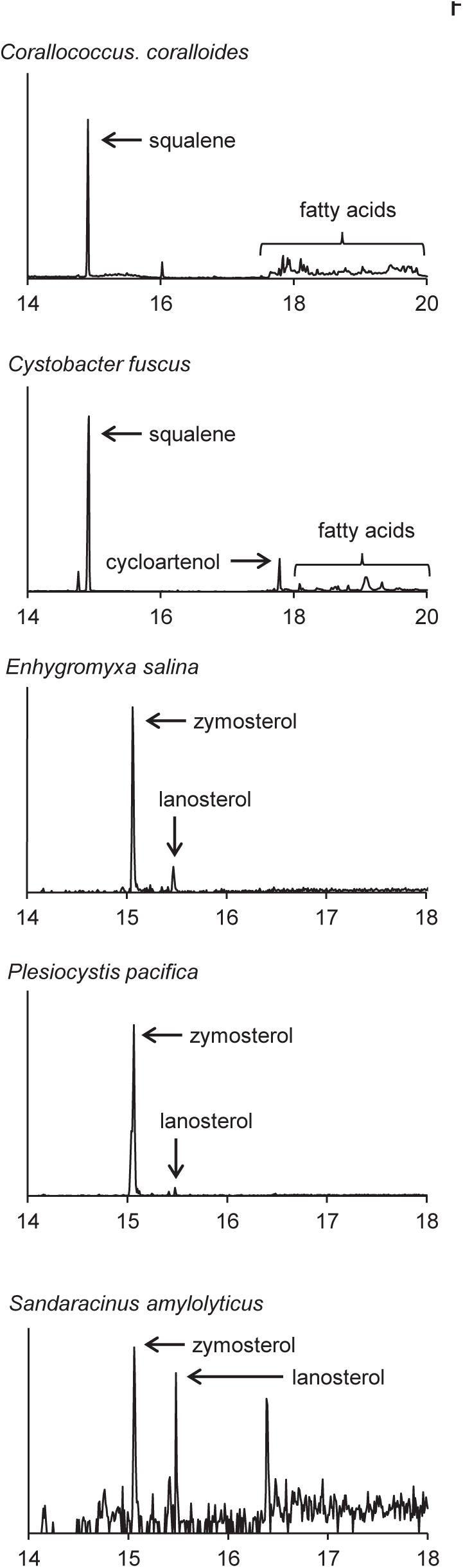
Sterols production in the myxobacteria. Extracted ion chromatograms (m/z 69, 440, 442, 454, 456, 468, and 498) of total lipid extract (TLE) from five myxobacteria. *C. coralloides* and *C. fuscus* TLEs were extracted from liquid cultures and were acetylated prior to running on the GC-MS. *E. salina*, *P. pacifica* and *S. amylolyticus* TLEs were extracted from cultures on a plate as growing them in liquid cultures was difficult. These TLEs were trimethylsilylated prior to running on the GC-MS. Sterol peaks were identified based on their mass spectra as shown in Figure 2.

**Table 3.** Sterols identified in bacterial strains

|  | squalene | cycloartenol | lanosterol | parkeol | 4,4-dimethylcholesta-8,24-dien-3-ol | 4,4-dimethylcholesta-8-en-3-ol | 4-methylcholesta-8,24-dien-3-ol | 4-methylcholesta-8-en-3-ol | zymosterol |
| --- | --- | --- | --- | --- | --- | --- | --- | --- | --- |
| **Myxobacteria** |  |  |  |  |  |  |  |  |  |
| *Corallococcus coralloides* | + |  |  |  |  |  |  |  |  |
| *Cystobacter fuscus* | + | + |  |  |  |  |  |  |  |
| *Enhygromyxa salina* |  |  | + |  |  |  |  |  | + |
| *Plesiocystis pacifica* |  |  | + |  |  |  |  |  | + |
| *Sandaracinus amylolyticus* |  |  | + |  |  |  |  |  | + |
| **Methylococcales** |  |  |  |  |  |  |  |  |  |
| *Methylobacter luteus* |  |  |  |  |  | + |  | + |  |
| *Methylobacter whittenburyi* |  |  |  |  | + | + | + | + |  |
| *Methylococcus capsulatus* Texas |  |  |  |  | + | + | + | + |  |
| *Methylosarcina lacus* |  |  |  |  | + |  | + |  |  |
| **Bacterioidetes** |  |  |  |  |  |  |  |  |  |
| *Fluviicola taffensis* |  | + |  |  |  |  |  |  |  |
| **α-Proteobacteria** |  |  |  |  |  |  |  |  |  |
| *Methyloceanibacter caenitepidi* |  | + |  |  |  |  |  |  |  |
| **Strains tested in previous studies** |  |  |  |  |  |  |  |  |  |
| *Nannocystis* species (Bode et al., 2003) | + |  | + |  | + | + | + | + | + |
| *Stigmatella aurantica* (Bode et al., 2003) | + | + |  |  |  |  |  |  |  |
| *Cystobacter* species (Bode et al., 2003) | + | + | + |  |  |  |  |  |  |
| *Corallococcus* species (Bode et al., 2003) | + |  |  |  |  |  |  |  |  |
| *Gemmata obscuriglobus* (Pearson et al., 2003) |  |  | + | + |  |  |  |  |  |
| *Methylococcus capsulatus* Bath (Bird et al., 1971;Bouvier et al., 1976) |  |  |  |  | + | + | + | + |  |
| *Methylomicrobium alcaliphilum* 20Z (Banta et al., 2015) |  |  |  |  |  | + |  |  |  |

**Table 4.** Identification of sterol biosynthesis genes in bacterial genomes

|  |  |  |  |  |  | C-4 demethylation |  |  |  |
| --- | --- | --- | --- | --- | --- | --- | --- | --- | --- |
|  | squalene oxygenation<br>SE | oxidosqualene<br>cyclization - Las<br>(lanosterol) | oxidosqualene<br>cyclization - Cas<br>(cycloartenol) | C-14 demethylation<br>ERG11 | C-14 reduction<br>ERG24 | C-4 methyl oxidation<br>ERG25 | C-3 dehydrogenation<br>ERG26 | C-3 ketoreduction<br>ERG27 | C-24(28) reduction<br>ERG4/DHCR24 |
| **Strains tested in this study** |  |  |  |  |  |  |  |  |  |
| *Coralloccoccus coralloides* | COCO<br>R_<br>01775 |  | COCO<br>R_<br>01777 | COCO<br>R_<br>02429 |  |  |  |  | COCOR<br>_<br>02182 |
| *Cystobacter fuscus* | D187_<br>003106 |  | D187_<br>003104 | D187_<br>005870 |  | D187_<br>005391 |  |  | D187_<br>001522 |
| *Enhygromyxa salina* | Ga0055<br>550_<br>102654 | Ga0055<br>550_<br>114516 |  | Ga0055<br>550_<br>103211 | Ga0055<br>550_<br>105926 | Ga0055<br>550_<br>101078 |  |  | Ga0055<br>550_<br>101115 |
| *Eudoraea adriatica* | G504D<br>RAF_<br>2315 | G504D<br>RAFT_<br>2316 |  |  |  |  |  |  |  |
| *Fluviicola taffensis* | Fluta_<br>3221 |  | Fluta_<br>3214 |  |  |  |  |  |  |
| *Methylobacter luteus* | MetluD<br>RAFT_<br>1256 | MetluD<br>RAFT_<br>1255 |  | MetluD<br>RAFT_<br>1253 |  |  |  |  | MetluD<br>RAFT_<br>1263 |
| *Methylobacter whittenburyi* | GY38D<br>RAFT_<br>3868 | GY38D<br>RAFT_<br>3867 |  | GY38D<br>RAFT_<br>3865 |  |  |  |  | GY38D<br>RAFT_<br>3875 |
| *Methyloceanibacter caenitepidi* | GL4_<br>3111 |  | GL4_<br>3110 |  |  |  |  |  |  |
| *Methylococcus capsulatus* Texas | H156D<br>RAF_<br>2531 | H156D<br>RAFT_<br>2530 |  | H156D<br>RAFT_<br>1746 |  |  |  |  | H156D<br>RAF_<br>0889 |
| *Methylosarcina lacus* | MetlaD<br>RAFT_<br>0846 | MetlaD<br>RAFT_<br>_0845 |  | MetlaD<br>RAFT_<br>_0843 |  |  |  |  |  |
| *Plesiocystis pacifica* | PPSIR1<br>_<br>02838 | PPSIR1<br>_02843 |  | PPSIR1<br>_<br>36894 | PPSIR<br>1_<br>23244 | PPSIR<br>1_<br>14275 | PPSIR1<br>_<br>17435 |  |  |
| *Sandaracinus amylolyticus* | Ga0055<br>546_<br>12562 | Ga0055<br>546_<br>18051 |  | Ga0055<br>546_<br>101121 | Ga0055<br>546_<br>17127 | Ga0055<br>546_<br>103226 | Ga0055<br>546_<br>17162 |  | Ga0055<br>546_<br>16346 |
| **Strains tested in previous studies** |  |  |  |  |  |  |  |  |  |
| *Nannocystis excedens* | Ga0008<br>035_<br>04851 | Ga0008<br>035_<br>04852 |  | Ga0008<br>035_<br>02418 | Ga0008<br>035_<br>01127 | Ga0008<br>035_<br>06338 | Ga0008<br>035_<br>00875 |  | Ga0008<br>035_<br>06674 |
| *Stigmatella aurantica* | STAUR<br>_<br>5420 |  | STAUR<br>_<br>5418 | STAUR<br>_<br>2030 |  | STAUR<br>_<br>2074 |  |  |  |
| *Gemmata obscuriglobus* | GobsU_<br>010100<br>006610 | GobsU_<br>010100<br>006605 |  |  |  |  |  |  |  |
| *Methylococcus*<br>*capsulatus* Bath | MCA<br>2872 | MCA<br>2873 |  | MCA<br>2711 |  |  |  |  | MCA<br>1404 |
| *Methylomicrobium*<br>*alcaliphilum* 20Z | MEAL<br>Z_<br>0767 | MEAL<br>Z_<br>0768 |  | MEAL<br>Z_<br>0770 |  |  |  |  | MEALZ<br>_<br>3890 |

In agreement with a previous study, the myxobacterium *C. coralloides* produces significant amounts of squalene but no sterol-like molecules despite having a copy of both squalene epoxidase (SE), required for the conversion of squalene to oxidosqualene prior to cyclization, and oxidosqualene cyclase in its genome (Table 4 and Figure 7) (Bode et al., 2003). To determine if the *C. coralloides* SE and Osc proteins were missing any necessary functional residues, we constructed an alignment of a subset of the bacterial SE and Osc homologs with four eukaryotic SE and Osc sequences (Figure 8). Both of these alignments indicate that key functional amino acid positions in the *C. coralloides* SE and Osc proteins are conserved and so the proteins are likely to be functional. It is also possible that the lack of sterol production may be due a lack of expression under the specific laboratory growth conditions we tested. Current studies are focused on growing *C. coralloides* under various conditions to induce sterol synthesis as well expressing the *C. coralloides* SE and Osc homologs in a heterologous system to verify that these proteins are functional.

**Figure 8.**
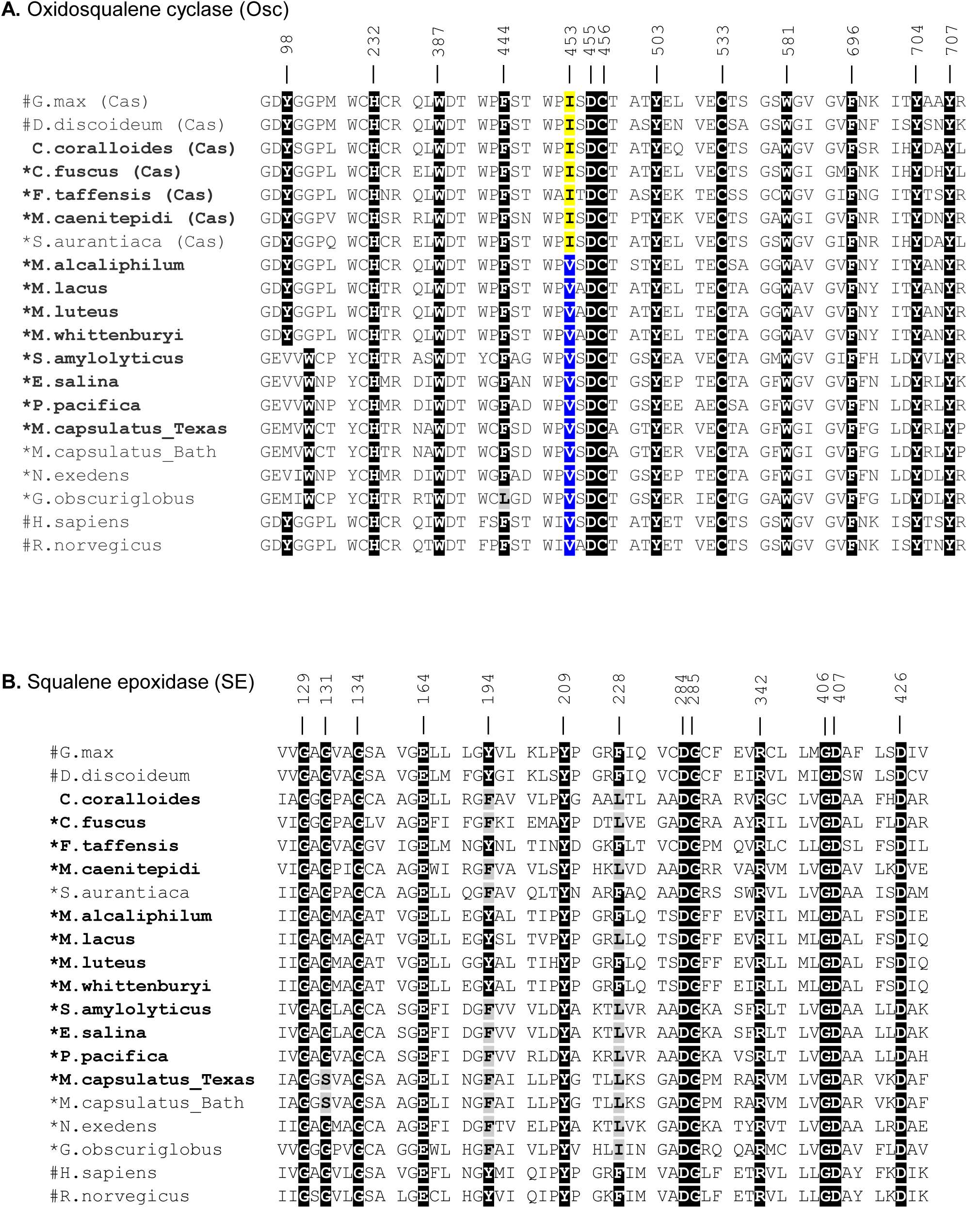
Amino acid alignments of the critical functional domains of oxidosqualene cyclase **(A)** and squalene epoxidase **(B)** homologs. Residues in black indicate residues that have been demonstrated to have a role in the biosynthesis of sterols in eukaryotes. Grey residues are those that differ from the conserved residue. In the Osc alignment, an isoleucine (I) at 453 (yellow) indicates a cycloartenol synthase and a valine (V) at 453 (blue) indicates a lanosterol synthase. Numbers correspond to residues in human Osc and SE. Bold labels indicate bacterial strains tested in this study. #: eukaryotic sequences, *: bacteria that have been shown to produce sterols.

### Sterol production in the methanotrophs

The lipid profiles of the four Methylococcales species tested were similar to what was previously observed in *M. capsulatus* Bath (Volkman, 2005), with some exceptions (Figure 9). *M. lacus* did not saturate the sterol side chain at C-24 as would be predicted because it lacks a homolog of the C-24(28) sterol reductase (ERG4 in yeast or DHCR24 in humans) (Table 4). *M. luteus*, on the other hand, only produced sterols that were saturated at the C-24 position (Figure 9). Interestingly, while all of the Methylococcales tested produced sterols that were partially demethylated at the C-4 position, none had homologs of any of the eukaryotic C-4 demethylase genes (Table 4). These methanotrophs also had sterols in which the unsaturation generated during C-14 demethylation was subsequently removed even though they lack a homolog of the C-14 reductase (ERG24) (Table 4). This is in contrast to the previously tested *M. alcaliphilum* (Banta et al., 2015) which does have a homolog of the C-14 reductase (ERG24) indicating that there may be more than one mechanism for this reaction even within the Methylococcales.

**Figure 9.**
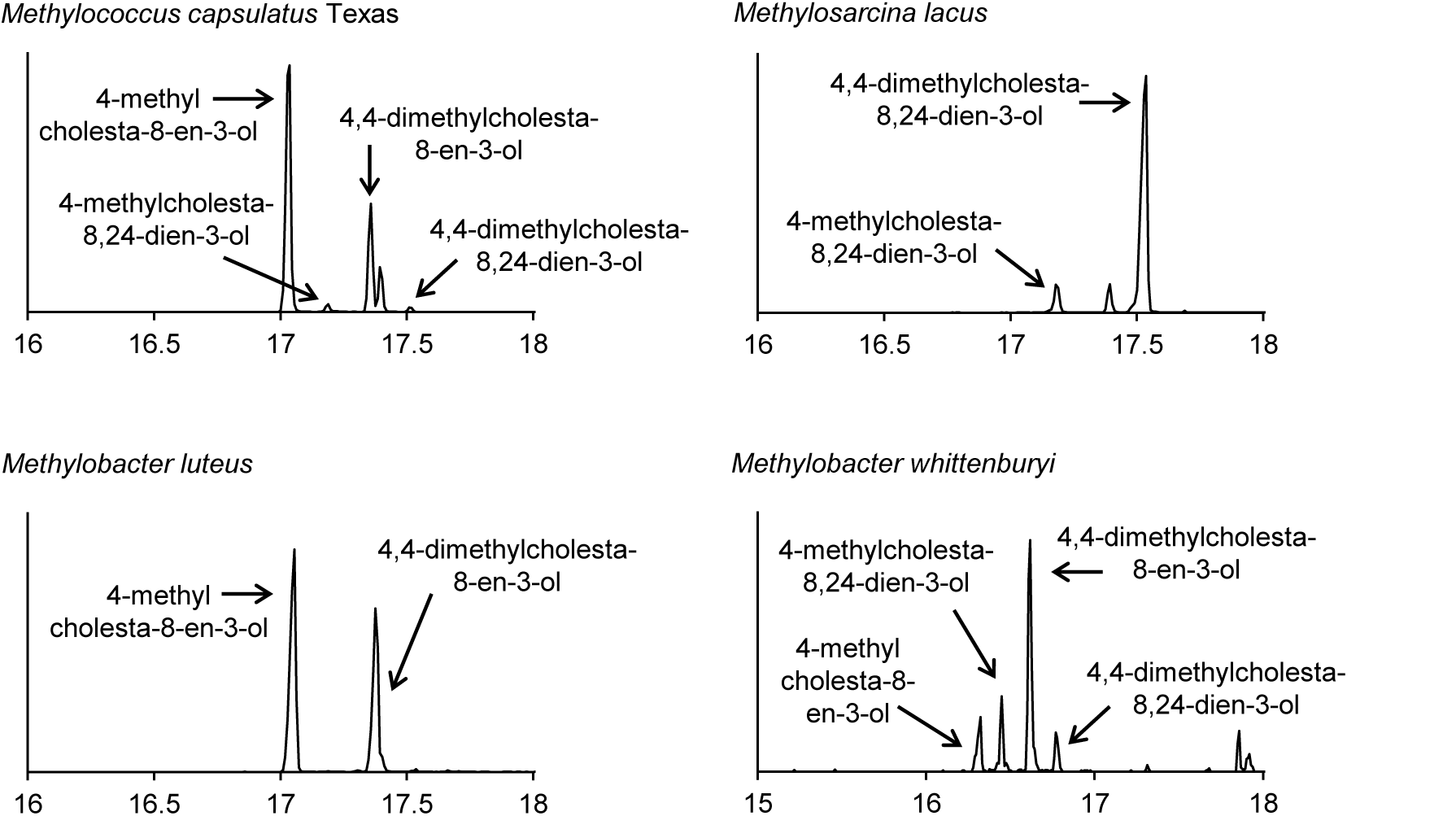
Sterols production in the aerobic methanotrophs. Extracted ion chromatograms (m/z 69, 440, 442, 454, 456, 468, and 498) of total lipid extract (TLE) from four aerobic methanotrophs. All TLEs were extracted from liquid cultures and were acetylated prior to running on the GC-MS. Sterol peaks were identified based on their mass spectra as shown in Figure 2.

### Sterol production in other bacterial species

We also observed production of cycloartenol in one Bacteriodetes species, *F. taffensis*, and one α-Protebacterium, *M. caenitepidi* (Figure 10 and Table 3). Neither of these strains had homologs of sterol biosynthesis genes downstream of *osc* in their genomes and this was in agreement with our observations of only cycloartenol production (Table 4).

**Figure 10.**
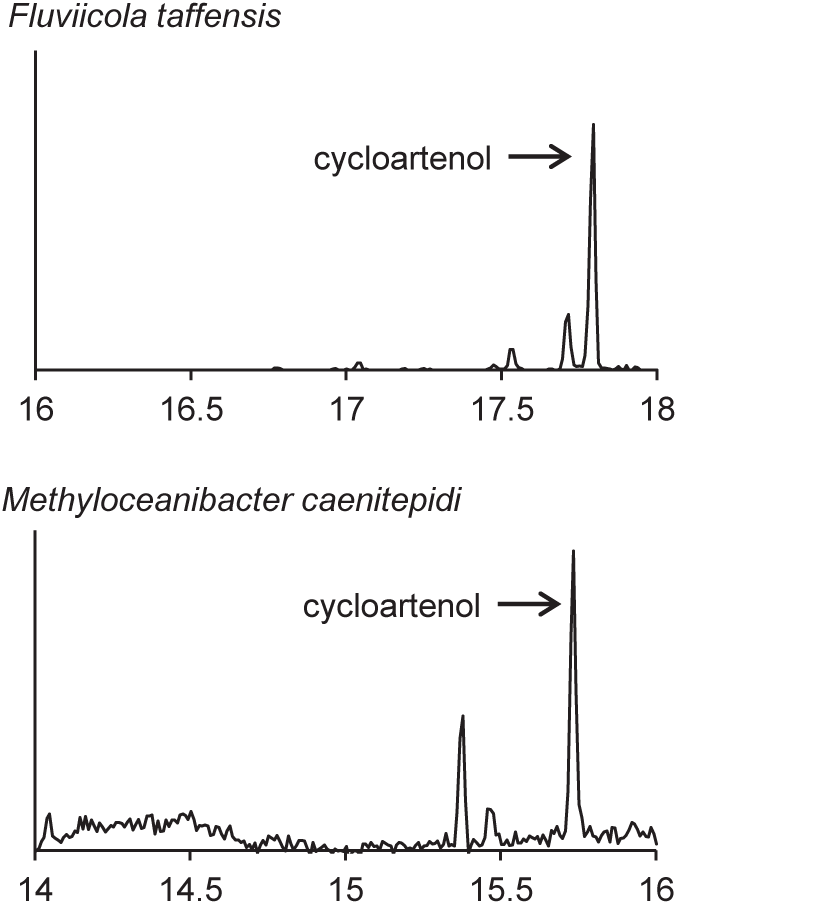
Sterols production in one Bacteriodetes and one α-Proteobacterium. Extracted ion chromatograms (m/z 69, 440, 442, 454, 456, 468, and 498) of total lipid extract (TLE) from the Bacteriodetes strains *F. taffensis* and the α-Proteobacterium *M. caenitepidi*. All TLEs were extracted from liquid cultures. The F. taffensis TLE was acetylated prior to running on the GC-MS. *M. caenitepidi* TLEs were trimethylsilylated prior to running on the GC-MS. Sterol peaks were identified based on their mass spectra as shown in Figure 2.

### Cycloartenol versus lanosterol synthesis is likely correlated with a single residue

The production of cycloartenol by some strains in our survey and lanosterol by others prompted us to investigate if specific residues were indicative of whether a cyclase was a lanosterol or cycloartenol synthase. Site-directed mutagenesis studies have previously identified three amino acids changes that seem to control the product profile of oxidosqualene cyclases (Meyer et al., 2000;Meyer et al., 2002;Lodeiro et al., 2004). Specifically, the amino acid residues T381/C,Q449/V453 (numbering based on human Osc) were indicative of a lanosterol synthase while Y381/H449/I453 suggested a cycloartenol synthase (Summons et al., 2006). Comparative genomics of three bacterial cyclases with eukaryotic cyclases revealed that only one of these residues was conserved and suggested that a valine (V) or isoleucine (I) at residue 453 suggested lanosterol or cycloartenol production, respectively (Summons et al., 2006). Our lipid analyses and alignments (Figure 8) verify that the bacterial oxidosqualene cyclases in the organisms we tested completely correlated with the observation that a V453 was indicative of lanosterol production while I453 signified cycloartenol production.

## Discussion

Sterol biosynthesis is primarily viewed as a eukaryotic feature that is rarely observed in the bacterial domain. Here, we coupled bioinformatics with lipid analyses to demonstrate that sterol production occurs in diverse bacteria and that this pathway may exist in yet to be discovered bacterial species. Our phylogenetic analysis of one of the key proteins involved in sterol biosynthesis, the oxidosqualene cyclase (Osc), demonstrates that the evolutionary history of this pathway in the bacterial domain is complex. In two previous phylogenomic studies it was concluded that sterol biosynthesis in the bacterial domain was most likely acquired through horizontal gene transfer (Desmond and Gribaldo, 2009;Frickey and Kannenberg, 2009). However, these phylogenetic analyses were limited as only three bacterial Osc sequences were available at the time these studies were undertaken. Our phylogenetic reconstruction with a larger data set demonstrates that bacterial Osc homologs fall into two clades – one monophyletic with the eukaryotic Osc sequences (Group 2) and one forming a sister clade to the eukaryotic sequences (Group 1). This topology implies that Group 2 *osc* genes were most likely acquired via horizontal transfer from a eukaryote as previously proposed (Frickey and Kannenberg, 2009) while Group 1 *osc* genes are more divergent from those found in eukaryotes. It seems plausible that Group 1 Osc sequences may represent a primitive lineage of cyclases suggesting that these bacteria may contain a more ancestral sterol biosynthetic pathway. However, more bacterial and eukaryotic Osc sequences as well as more rigorous phylogenetic analyses are needed to better interpret the evolutionary history of sterol biosynthesis in both the bacterial and eukaryotic domains.

While our phylogenetic analyses suggest a complex evolutionary history of sterol biosynthesis in bacteria, our lipid analyses demonstrate less modification of sterols in bacteria compared to what is usually observed in eukaryotes. Many of the bacteria we tested produced lanosterol or cycloartenol as the end product. Production of these basic sterols only requires two biosynthetic steps of the canonical eukaryotic sterol pathway - the epoxidation of squalene to oxidosqualene and the subsequent cyclization of oxidosqualene to lanosterol or cycloartenol. The myxobacteria and methanotrophs, however, did make certain modifications such as C-4 and C-14 demethylations and isomerization of double bonds in the main ring structure. Interestingly, not all proteins required to make those modification in eukaryotes were found in the genomes of these bacteria. In particular, the removal of the C-4 methyl groups requires the activity of three eukaryotic proteins, a C-4 methyl oxidase (ERG25), a C-4 decarboxylase (ERG26) and a C-3 ketoreducatse (ERG27) (Bard et al., 1996;Gachotte et al., 1998;Gachotte et al., 1999). These three proteins were first identified in yeast and homologs have been identified in most sterol producing eukaryotic genomes, with the exception of plants which seem to be missing an ERG27 homolog (Desmond and Gribaldo, 2009). In the myxobacteria, we observed that two of the organisms that removed the C-4 methyl groups had homologs of ERG25 and ERG26 but not ERG27 and a third organism only had a homolog of ERG25. Desmond and Gribaldo attempted to identify potential ERG27 homologs in the genome of *P. pacifica* through comparative genomics. One potential gene candidate was identified (*P. pacifica* locus tag: Ga0067453_11974) and the myxobacteria we tested do have a homolog of this protein in their genomes. However, further studies are needed to determine if this protein is necessary for C-4 demethylation in the myxobacteria.

It is also possible that downstream sterol modifications in bacteria occur via distinct biochemical pathways than what is observed in eukaryotes. This is a particularly compelling in the aerobic methanotrophs. In these organisms, one methyl group is removed at the C-4 position but we could not identify homologs of the eukaryotic C-4 demethylase genes (ERG25, ERG26 or ERG27). In addition, we observed saturation of the C-14 double bond in the sterols of all methanotrophs tested but did not identify a C-14 reductase (ERG24) in their genomes. The discrepancies in the sterols produced by methanotrophs and the proteins identified in their genomes points to the possibility that novel sterol biosynthesis proteins may exist in bacteria. Identification and characterization of these bacterial sterol proteins could reveal unique biochemical and regulatory mechanisms. In addition, a full understanding of the proteins involved in bacterial sterol production will allow for studies to discern what functional role these lipids play in the bacterial cell and would provide significant insight into the evolution of this ancient biosynthetic pathway.

Finally, this work demonstrates the utility of combining bioinformatics with lipid analyses to get a broader picture of not just sterol synthesis in bacteria but potentially other geologically relevant lipids. The increasing amount of genome and metagenome sequence data available along with advancements in developing culturing and genetic systems in nontraditional microbes provides an excellent opportunity for exploring many aspects of biomarker lipids in microbes – their biosynthesis, their function and their evolutionary history. Ultimately, a full understanding of microbial biomarker lipids will provide valuable information for more nuanced interpretation of microbial lipid biosignatures in both modern ecosystems and ancient sedimentary rocks.

## Author contributions

PW designed the study. JW, XY and PW acquired the data. JW and PW analyzed the data. PW wrote the manuscript.

## Acknowledgments

We would like to thank Prof. Marina G. Kalyuzhnaya for her contribution of several of the methanotroph strains utilized in this study. We would also like to thank Dr. Amy B. Banta and other members of the Welander research group for thoughtful discussions of this work and helpful comments on the manuscript.

